# Loss of *prdm1a* accelerates melanoma onset and progression

**DOI:** 10.1101/2019.12.20.884767

**Authors:** Ritsuko Iwanaga, Brittany T. Truong, Jessica Y. Hsu, Karoline A. Lambert, Rajesh Vyas, David Orlicky, Yiqun G. Shellman, Aik-Choon Tan, Craig Ceol, Kristin Bruk Artinger

## Abstract

Melanoma is an aggressive, deadly skin cancer derived from melanocytes, a neural crest cell derivative. Melanoma cells mirror the developmental program of neural crest cells in that they exhibit the same gene expression patterns and utilize similar cellular mechanisms, including increased cell proliferation, EMT and migration. Here we studied the role of neural crest regulator PRDM1 in melanoma onset and progression. In development, Prdm1a functions to promote neural crest progenitor fate, and in melanoma, we found that *PRDM1* has reduced copy number and is recurrently deleted in both zebrafish and humans. When examining expression of neural crest and melanocyte development genes, we show that *sox10* progenitor expression is high in *prdm1a-/-* mutants, while more differentiated melanocyte markers are reduced, suggesting that normally Prdm1a is required for differentiation. Data mining of human melanoma datasets indicate that high *PRDM1* expression in human melanoma is correlated with better patient survival and decreased *PRDM1* expression is common in metastatic tumors. When one copy of *prdm1a* is lost in the zebrafish melanoma model (Tg[mitfa:BRAF^V600E^];p53-/-; *prdm1a*+ /-), melanoma onset occurs more quickly, and the tumors that form have a larger area with increased expression of *sox10.* These data demonstrate a novel role for PRDM1 as a tumor suppressor in melanoma.

## INTRODUCTION

Melanoma is an aggressive cancer that accounts for the majority of human skin cancer deaths. The 5-year survival rate of metastatic melanoma is only 23% ^1^. Early stage melanoma can be cured by surgical excision; however, only a few treatment options exist once the melanoma has metastasized. More than 50%of melanoma patients present with a somatic *BRAF* mutation^2^, and up to 90%of these are from a single amino acid change from valine (V) to glutamic acid (E) at position 600 (V600E)^3^. As such, the high frequency of *BRAFV600E* mutations among melanoma patients has prompted the development of targeted therapies such as BRAF and MEK inhibitors that dampen BRAF/MEK pathway signaling^4^. Dabrafenib and trametinib were the first combined inhibitors to be approved by the Food and Drug Administration (FDA) for patients with advanced *BRAFV600E* melanoma^5,6^. The combined inhibitors improve patient survival rates and have fewer adverse side effects compared to monotherapy treatments. About one third of melanoma patients experience disease recurrence or metastasis years after initial treatment^7,8^. The acquirement of additional somatic mutations (i.e. point mutations, chromosomal rearrangements, copy number variations, etc.) that bypass the targets of BRAF and MEK inhibitors may account for some cases of incomplete response to the primary line of targeted therapy. However, other mechanisms of resistance or melanoma progression exist that have yet to be studied. For example, we previously showed that gene amplification of *SETDB1*, in conjunction with a *BRAF*^*V600E*^ mutation, accelerates melanoma tumor formation in a zebrafish model by causing disruptions in transcriptional regulation, including a decrease in expression of *HOX* genes^9^. Identifying additional genes and pathways involved in melanoma initiation is critical, as it can lead to the development of future therapeutic and treatment options.

The cell origins of melanoma tumors are thought to arise from transformed melanocytes^10^, which are derived from neural crest cells, a multipotent population of cells that forms between the neural and non-neural ectoderm. During embryonic development, neural crest cells undergo an epithelial-to-mesenchymal transition and migrate to different regions of the developing embryo before giving rise to a variety of cell types, including peripheral nerves, chondrocytes, osteoblasts, and melanocytes. As neural crest cells are a highly migratory population of cells, it has been hypothesized that the same genes and pathways utilized by neural crest cells during development to aid in the trafficking of cells to their final destinations in the embryo can be later hijacked by cancer cells to enhance their proliferation and migration^11,12^. Indeed, several publications have shown that there are distinct parallels in behaviors and gene expression patterns between melanoma cells and neural crest cells^13^. For instance, Brn3a, a neural crest transcription factor in the neuronal lineage, is frequently expressed in melanoma tumors and has been shown to accelerate tumor growth by promoting survival^14^. Interestingly, mouse melanoma cells can arise from a population of seemingly differentiated melanocytes that respond to environmental cues to undergo transcriptional reprograming^15^. One cue is likely a neural crest growth factor, as melanoma cells express CD271 receptors on the cell surface^16^.

Apart from these studies that have implicated single neural crest-related genes in melanoma progression, there is strong evidence that a neural crest transcriptional program plays a causative role in melanoma progression. Genes such as *crestin, sox10, ednrb, tyr*, and *dct*, which are characteristic markers of neural crest progenitors and melanocytes, are highly enriched in zebrafish and mouse melanoma tumors^17,18^. In zebrafish, *crestin* is normally downregulated at later stages in development after neural crest cells have fully differentiated^19^. Using a *crestin:EGFP* reporter in a zebrafish model, Kaufman et al. (2016) used live-cell imaging of melanoma tumor formation and showed that a single melanocyte can initiate tumor growth by reactivating *crestin*^20^. Moreover, *crestin:EGFP-*positive zebrafish scales, but not adjacent *crestin-*negative scales, exhibit gene signatures that were enriched with neural crest and melanoma genes, such as *mitfa* and *sox10*^20^. This work suggests that reactivation of specific neural crest cell genes coincides with tumor initiation. It is evident that inappropriate activation of developmental genes can lead to cancer; however, it is not well-known whether expression of neural crest cell genes can also be protective.

Here, we identify neural crest regulator PRDM1 as a novel tumor suppressor in melanoma. *PRDM1* (zebrafish ortholog *prdm1a*) is a highly conserved transcription factor, whose structure consists of an N-terminal PR/SET domain, followed by a proline-rich domain, and five DNA-binding C2H2 zinc fingers. PRDM1 regulates transcription both by direct DNA binding^21^ and by forming complexes with chromatin-modifying proteins including histone methyltransferases Prmt5^22^ and G9a^23^ and histone deacetylases HDAC1/2^24^. *PRDM1* is a critical factor in neural crest cell development. It is highly expressed at the neural plate border in zebrafish^25,26^, *Xenopus laevis*^27^, and lamprey^28^ and is required for Rohon-Beard sensory neuron and neural crest cell formation^25,26^. Homozygous loss of *prdm1a* in developing zebrafish embryos results in a significant reduction of neural crest cell derivatives, including pigment cells and craniofacial cartilage^25^, suggesting that it is required for neural crest differentiation. Similarly, PRDM1 has also been shown to be required for the differentiation of both adaxial cells into slow-twitch muscle cells^29^, as well as retinal progenitors to photoreceptor cells^30,31^. In addition to being important for cell fate specification during development, PRDM1 and other members of the PRDM family function as tumor suppressors for many cancers, including B, T, and natural killer cell lymphomas, colorectal cancer, and lung cancer^32-35^. In lymphoid malignancies, genomic deletions, missense mutations, and premature truncations of *PRDM1* are frequently observed^34,36,37^, causing inactivation of the protein and misregulation of lymphocyte differentiation and cell cycle progression. These studies provide strong evidence and support a role for PRDM1 as a tumor suppressor. In this study, we show that Prdm1a is required for melanocyte differentiation from neural crest cells during development and that decreased *PRDM1* expression is correlated with worse melanoma patient survival. We also report that loss of *PRDM1* in a zebrafish model leads to an acceleration of melanoma onset and progression.

## MATERIALS & METHODS

### Zebrafish

Zebrafish were maintained as previously described^38^. The Institutional Animal Care and Use Committee (IACUC) at the University of Colorado Denver Anschutz Medical Campus (UCD-AMC) reviewed and approved all animal procedures. Wildtype strains include AB and EKK lines. Transgenic lines include *prdm1a*^*m805*^ *(nrd)*^*39*^, referred to as *prdm1a-/-*; *tp53*^*M214/M214*^, referred to as *p53-/-*; Tg(*mitfa*:BRAF^V600E^);p53-/-^17,40^, and Tg(*mitfa*:BRAF^V600E^);p53-/-.

The standard length of the zebrafish was measured from the tip of the snout to the base of the tail (**Figure 2A**).

**Figure 1:**
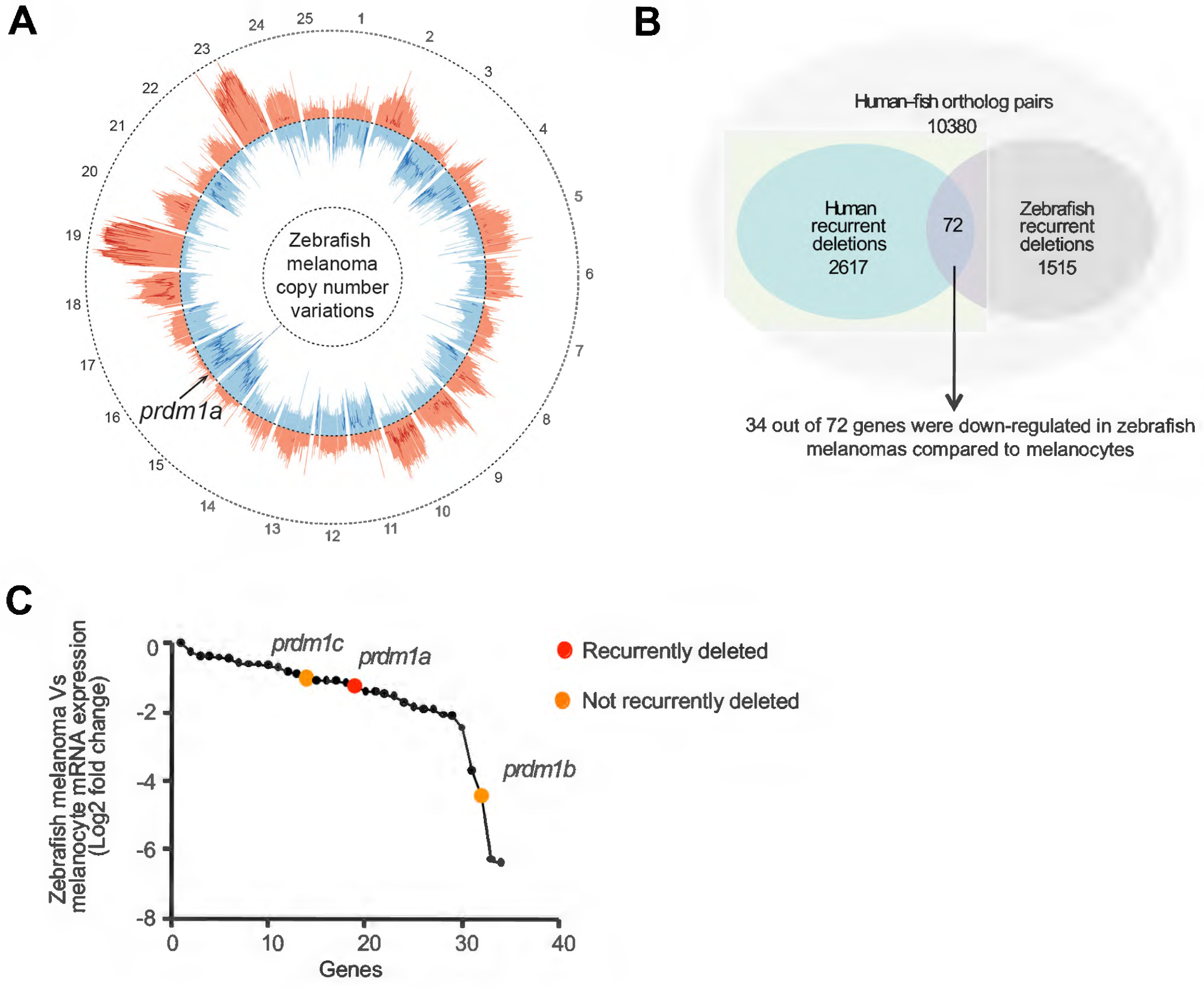
PRDM1 is recurrently deleted in human and zebrafish melanoma. (A) Circos plot displaying gene copy number gains and losses in zebrafish melanomas as previously described^41^. JISTIC G-scores are displayed as pale red shading (amplifications [minimum = 0; maximum = 1,550]) and blue shading (deletions [minimum = 0; maximum = 2,150]). –log10-transformed JISTIC Q-values with a cutoff of 0.6 (corresponding to an untransformed Q-value of 0.25) are shown as bold red lines (amplifications [minimum = 0; maximum = 11]) and bold blue (deletions [minimum = 0; maximum = 11]). Dotted circles represent the –log10-transformed Q-value of 0 (center) and 11 (outer: amplification; inner: deletion). (B) Venn diagram of orthologous genes significantly deleted in human and zebrafish melanomas from 10,380 human-zebrafish gene pairs (hypergeometric test, p-value: 2.0E-16). (C) Genes downregulated in zebrafish melanomas as compared with melanocytes (analyzed from previously obtained microarray data set^41^). *prdm1a* (red dot) and its paralogs *prdm1b* and *prdm1c* (orange dots) are indicated.

**Figure 2:**
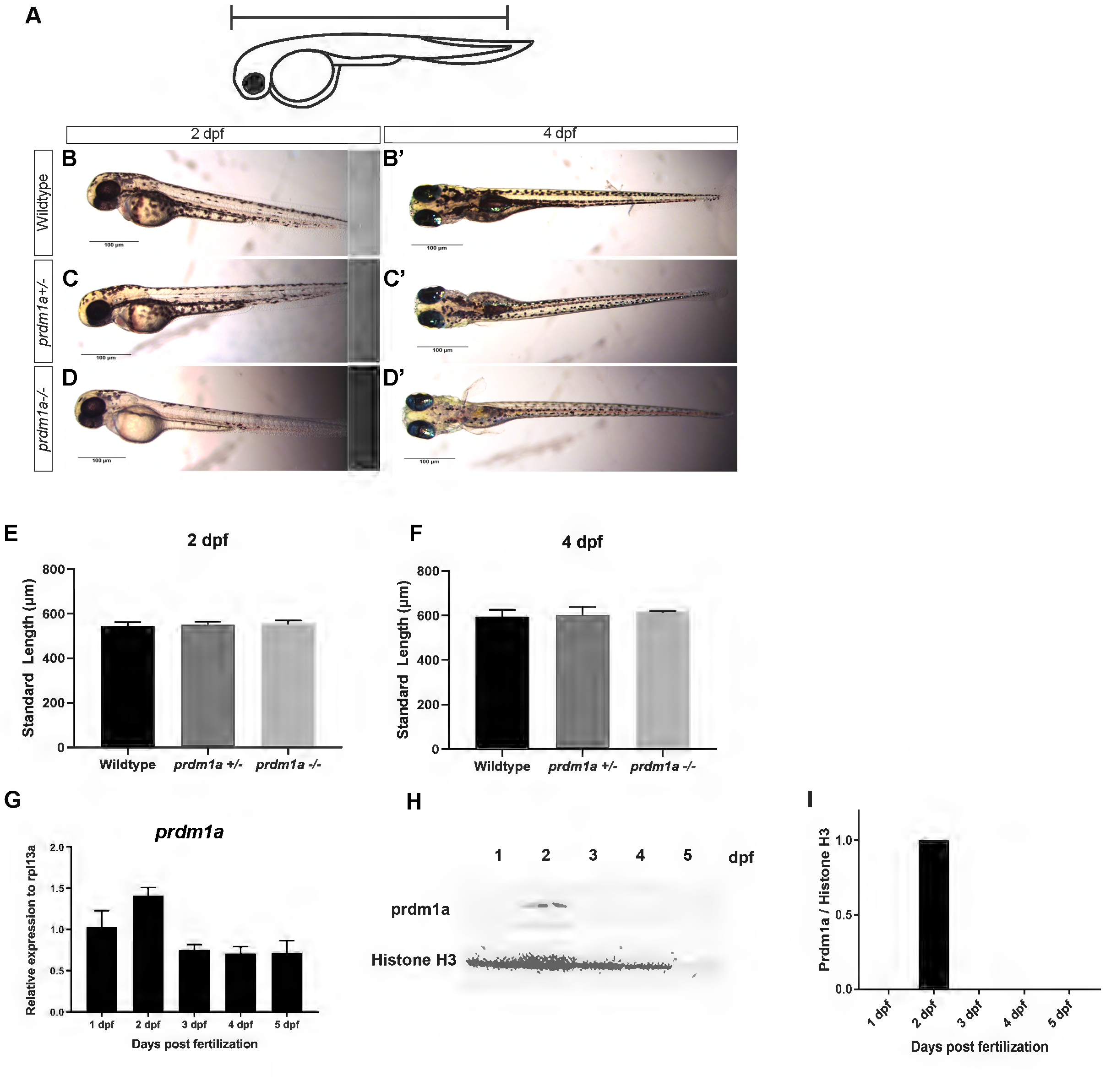
prdm1a. (A) Schematic showing how the standard length of the embryo was measured. Lateral images of wildtype (B), *prdm1a+ /-* (C) and *prdm1a-/-* (D) at 2 dpf. Ventral images of wildtype (B’), *prdm1a+ /-* (C’) and *prdm1a-/-* (D’) at 4 dpf. Quantification of standard length at 2 (E) and 4 (F) dpf. Statistical analysis was performed with a one-way ANOVA. (G) RT-qPCR was performed on 20 pooled embryos at each time point (n=3 replicate experiments) for *prdm1a*. Proteins were resolved by SDS-PAGE and western blotting for *prdm1a* and total H3 (H) was performed. (I) Quantification of band intensity from the western blot (G). Relative intensity was normalized to total histone H3 for each sample. Abbreviations: dpf, days post fertilization

Microinjections were performed in p53-/- and *prdm1a+/-* zebrafish at the single-cell stage. The *mitfa:*BRAF^V600E^ vector was a gift from Richard M. White^17^. The vector was linearized with SalI and BssHI, purified, and resuspended in 100 mM KCl for a final concentration of 100 ng/mL. 500 pL were injected into each embryo.

Zebrafish were observed and scored weekly for hyperpigmentation, which indicated the presence of melanoma tumors. We compared tumor-free curves over time using a log-rank test where p< 0.05 was statistically significant. The tumor area was measured using ImageJ and normalized to body length.

### aCGH, JISTIC, and comparative analysis

Array comparative genomic hybridization (aCGH) was performed and analyzed as previously described^41^. Briefly, we custom designed an array with 398,426 unique probes covering 97%of the zebrafish genome (based on Zv9 assembly). Genomic DNA was extracted from zebrafish melanoma tumors (Tg[*mitfa*:BRAF^V600E^];p53-/- ^17,40^) and from a normal region of the same fish using the Qiagen DNeasy Blood and Tissue kit (Qiagen, Germantown, MD, #69504). The DNA was fragmented and hybridized to the array. Raw data was generated from scanned images with the Agilent Feature Extraction software (v10.7) and normalized using the Agilent Genomic workbench to identify copy number differences (Agilent, Santa Clara, CA). The JISTIC algorithm was used to calculate significantly altered regions, and peak calling was done using a q-value cut-off of 0.25. For human melanomas, copy number data was downloaded from Tumorscape^42,43^ and JISTIC analysis was conducted as described previously^41^. Genes from within peaks were pooled to define species-specific sets of recurrently deleted genes. Human orthologs of zebrafish genes were determined using Ensembl and BLAST.

### Survival analysis of The Cancer Genome Atlas (TCGA) cutaneous melanoma data

Clinical and mutational data of 329 TCGA melanoma patients were obtained from the supplementary tables of the TCGA publication (http://cancergenome.nih.gov/). Normalized RNA-seq gene expression profiles (Level 3, RSEM value) for these patients were downloaded from the cBioPortal website (http://www.cbioportal.org). Patients were assigned to high (hi) and low (lo) groups based on above or below mean gene expression of *PRDM1*. In this study, we only focus on the 286 metastatic melanoma patients with outcome data. Kaplan-Meier survival analysis was performed comparing hi and lo groups. A log-rank test of p<0.05 was considered statistically significant.

### Mining public gene expression data sets

Gene expression profiles of the Riker data set (GSE7553)^44^ were downloaded from the Gene Expression Omnibus (GEO) (http://www.ncbi.nlm.nih.gov/gds). This data set contains 87 melanoma patients and was profiled by the Affymetrix Human Genome U133 Plus 2.0 microarray. Raw data was downloaded and normalized with the Robust Multiarray Average (RMA) method using the Affymetrix Power Tools. From this data set, there were 4 normal skin, 14 primary melanoma and 40 metastatic melanoma patients. Gene expression for *PRDM1* was compared across these patients, and a one-way ANOVA test was performed to determine the different expressions. Error bars represent mean ± SD.

### Histology

Zebrafish samples were fixed in 4%paraformaldehyde and embedded in paraffin. The samples were then sectioned at a 5 μM thickness and fixed to glass slides. H&E staining was performed by the Gates Center Regenerative Medicine Histology Core at UCD-AMC. Histology images were captured on an Olympus BX51 microscope equipped with a four megapixel Macrofire digital camera (Optronics, Goleta, CA) using the Picture Frame Application 2.3 (Optronics).

### RNA extraction and real-time PCR (RT-qPCR)

Total RNA was isolated from dissected zebrafish tumors and zebrafish embryos (20 pooled embryos). The zebrafish samples were stored in TRIzol reagent (Invitrogen Corp., Carlsbad, CA, #15596018) and then purified with the Direct-zol RNA kit (Zymo Research, Irvine, CA). RNA (1 μg) was converted to cDNA using the Verso cDNA Synthesis Kit (Thermo Fisher Scientific, Waltham, MA). All procedures were conducted according to the manufacturers’ suggestions.

Real-time PCR reactions (RT-qPCR) were performed in the Step One Plus Real-time PCR System (Thermo Fisher Scientific). Taqman primers for *PRDM1, prdm1a, sox10, dct, mitfa, mc1r*, and *tyr* were purchased from Thermo Fisher Scientific. *rpl13a* was used as the zebrafish internal control. (*rpl13a:* probe 5’-/56-FAM/ATGAAGTTGGCTGGAAGTACCA/3lABlk_FQ/- 3’, forward 5’-CACACGCAAATTTGCCCTGCTTGG-3’, and reverse 5’-GCCTGCTTAGTCAGCTTCATC-3’). Reactions were performed in three biological and technical replicates. Transcript abundance and relative fold change were quantified using the 2^-ΔΔCt^ method relative to control. Relative abundance was compared using an unpaired, independent t-test. Error bars represent the mean ± SD. Results were statistically significant when p< 0.05.

### Western blotting

Western blotting was performed as previously described^45^. 50 to 100 zebrafish embryos were pooled at each time point. Calcium-free Ginzberg Fish Ringer’s solution (NaCl, KCl, NaHCO3) was added to the embryos for deyolking. Samples were then washed in deyolking wash buffer (NaCl, Tris [pH 8.5], KCl, and CaCl2) and lysed in lysis buffer using a syringe (0.1%glycerol, 0.01% SDS, 0.1M Tris-HCL [pH6.8]). Total protein concentration was determined with the Bio-Rad DC Protein assay (Bio-Rad, Hercules, CA). Proteins (20 μg) were separated by SDS-PAGE (12%) and transferred to polyvinylidene difluoride membranes. Primary antibodies against PRDM1 (1:750 dilution, gift from Dr. Philip W. Ingham)^46^ and total histone H3 (1:5000 dilution, Cell Signaling Technology, Danvers, MA, #9715) were used. The blots were incubated with secondary antibody, HRP-conjugated goat anti-rabbit (1:5000 dilution, Sigma-Aldrich, St. Louis, MO, #A0545) for 2 hours. Chemiluminescent detection was performed with Luminata Classico Western HRP substrate (Millipore Sigma, Burlington, MA). Bio-Rad Chemidoc multiplex imaging system was used to detect and quantify the protein levels. Band intensity was quantified in ImageJ and normalized to the total histone H3 loading control. The experiment was performed at least twice.

### Tyrosinase assay

The tyrosinase assay was performed as previously described with a few modifications^47,48^. About 100 embryos at two days post fertilization (dpf) were dechorionated and sonicated in Pro-Prep protein extraction solution (iNtRON Biotechnology, Burlington, MA, #17081). 200 μg of total protein in 75 uL of lysis buffer was added to 75 μL of 1.0 mM of L-3,4-dihydroxyphenylalanine (L-DOPA) in a 96-well plate. The control well contained 75 μL of lysis buffer and 75 μL of 1.0 mM L-DOPA. After incubation at 37°C for 60 minutes, absorbance was read at 425 nm. The assay was performed twice with three technical replicates in each trial. Error bars represent the mean ± SD. The data were analyzed using a one-way ANOVA test followed by a multiple comparisons test where p< 0.05 was statistically significant.

## RESULTS

### PRDM1 is recurrently deleted in human and zebrafish melanoma and is required for melanocyte differentiation

Somatic gene mutations, including gene copy number variations, are frequently observed in cancer, including melanoma^9,43,49,50^. We used a cross-species oncogenomics approach to identify recurrently deleted genes in both human and zebrafish melanomas. As previously described, we performed aCGH on Tg(*mitfa*:BRAF^V600E^);p53-/- zebrafish to generate copy number variant (CNV) profiles for zebrafish that develop melanomas (see below)^17,40,41^ (**Figure 1A**). Human data were obtained from Tumorscape, a dataset that contains copy-number analyses for over 3,000 human cancer specimens^42,43^. The aCGH values and human data were analyzed using the JISTIC algorithm^51^, which then listed recurrently amplified and deleted genes. We identified 2,617 genes recurrently deleted in human melanomas and 1,515 genes in zebrafish melanomas. Of these recurrently deleted genes, 72 were deleted in both human and zebrafish melanomas (**Figure 1B**), and furthermore 34 of these genes were also identified as transcriptionally downregulated in a microarray dataset comparing expression in zebrafish melanoma tissue and normal melanocytes^41^. Among the 34 downregulated genes in zebrafish melanoma was the family of *prdm1* paralogs, *prdm1a-c* (**Figure 1C**). *prdm1a* was selected as a candidate gene because of its known role as a neural crest regulator^25^. Paralogs *prdm1b* and *prdm1c* were also downregulated, but they were not recurrently deleted based on the JISTIC analysis (**Figure 1C**). Therefore, we did not explore these paralogs further.

PRDM1 is a known tumor suppressor in multiple cancers, including B and T-cell lymphoma^36,52^, colorectal cancer^53^, lung cancer^54^, and glioma^55^. Moreover, Prdm1a has been shown to play a role in neural crest differentiation in zebrafish^25,39^. Homozygous loss of *prdm1a* results in reduced neural crest cell derivatives, including peripheral neurons and mature pigment cells (**Figure 2A-D**)^39^. *prdm1a-/-* mutants die at 6-7 dpf, and wildtype and heterozygotes are indistinguishable by phenotype (**Figure 2B-C’**). The standard length of the zebrafish from the tip of the snout to the base of the tail is not affected when one or both copies of *prdm1a* are mutated (p>0.05) (**Figure 2E-F**). *prdm1a* transcripts (**Figure 2G**) and protein levels (**Figure 2H-I**) are highest a 2 days post fertilization (dpf), when pigment cells are differentiating along body of the zebrafish^56^. To better understand the role of Prdm1a in melanocyte development, we performed RT-qPCR on whole wildtype and *prdm1a -/-* mutant embryos for a series of genes that mark early (*sox10*) and late melanocyte lineage (*dct, mc1r, mitfa, tyr*)^10^ during a time course of development from 1-5 dpf (Schematic **Figure 3A**). When neural crest cells commit to the melanocyte lineage, they express different markers that indicate their stage of differentiation (**Figure 3A**). *sox10* is an early melanocyte stem cell marker and is required for melanocyte differentiation from neural crest cells in that it activates expression of *mitf, dct*, and *tyr*^10,57^. Interestingly, in wildtype embryos, *sox10* expression is steady over time, but *prdm1a-/-* mutants exhibit a significant increase in expression at 2 and 4 dpf, corresponding to the first and second waves of melanocyte migration during zebrafish development^58^ (p< 0.0001) (**Figure 3B)**. Expression of *dct, mitfa*, and *tyr*, markers of the melanoblast precursor state, decrease steadily over developmental time, while *mitfa* increases in wildtype embryos (**Figure 3C-F**). In contrast, the *prdm1a-/-* mutants consistently exhibit an overall decrease in the later melanocyte markers compared to wildtype embryos (p< 0.05) (**Figure 3C-F**). The increase in *sox10*, along with the overall decrease in later melanocyte markers, suggests that the cells could be maintained as early melanocyte progenitors, unable to fully differentiate into mature, pigment producing melanocytes.

**Figure 3:**
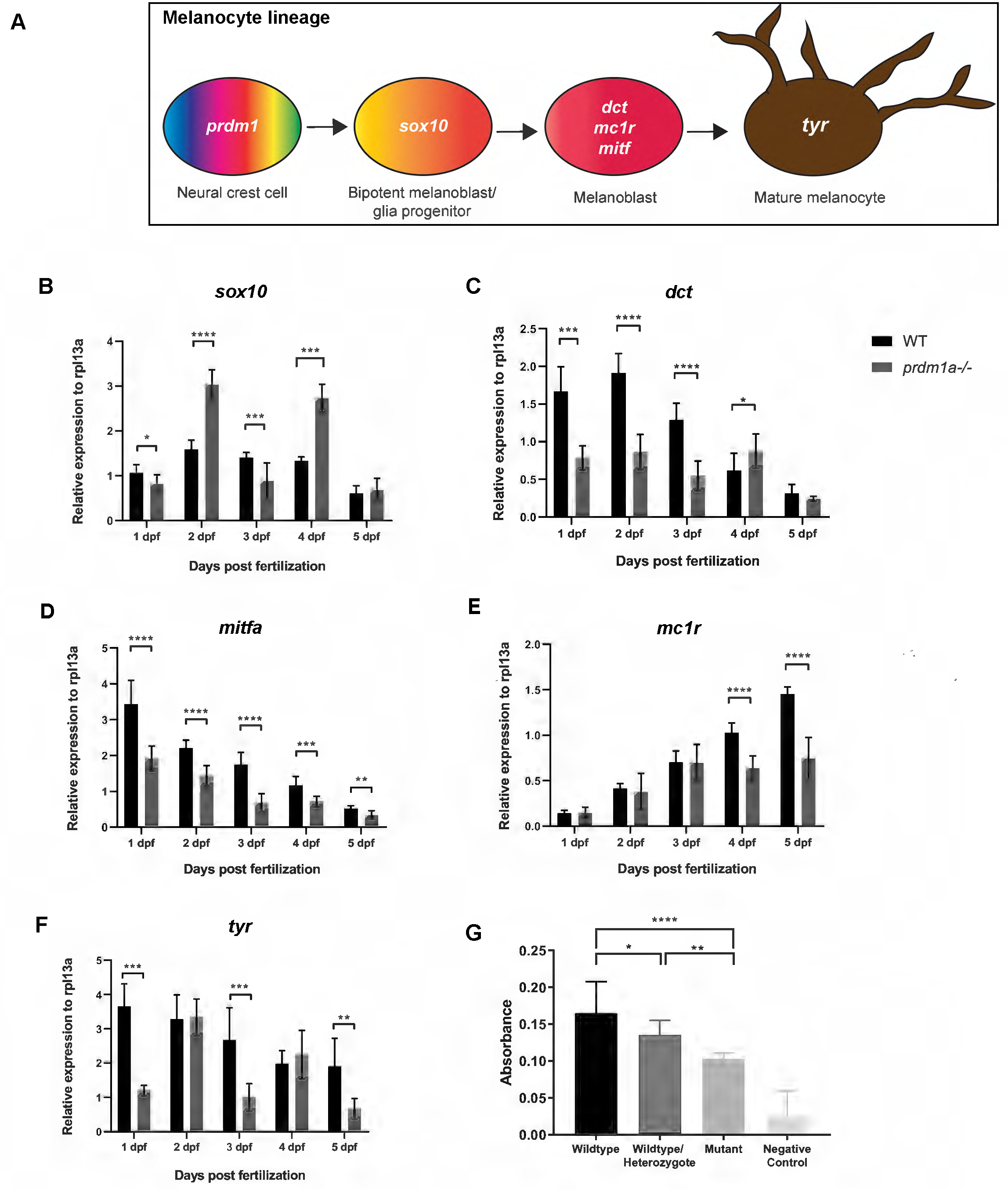
*prdm1a* is required for melanocyte differentiation and pigment production. (A) Melanocyte lineage from neural crest to mature melanocytes and their characteristic markers. Whole wildtype embryos were pooled for RNA isolation and protein extraction over developmental time. (B-F) Whole wildtype and *prdm1a-/-* mutant embryos were pooled for RNA isolation over developmental time. 20 whole embryos were pooled for each genotype and at each time point (n=3 replicate experiments). RT-qPCR was performed for *sox10* (B), *dct* (C), *mitfa* (D), *mc1r* (E), and *tyr* (F) over developmental time. Expression was compared using an unpaired, independent t-test. (J) 100 embryos were pooled and incubated with L-DOPA for a tyrosinase functional assay. Pigment production was measured by spectrometry in whole wildtype, wildtype/heterozygote mix, and *prdm1a-/-* mutant embryos. Statistical analysis was performed with a one-way ANOVA followed by a multiple comparisons test. Error bars represent mean ± SD. *p< 0.05, **p<0.01, ***p< 0.001, ****p< 0.0001. Abbreviations: dpf, days post fertilization

To further test whether loss of *prdm1a* affected proper melanogenesis, we tested for tyrosinase activity, the enzyme responsible for pigment production, in *prdm1a-/-* mutant, wildtype (EKK strain), and wildtype/heterozygote mixed embryos at 2 dpf. Pooled embryos were incubated with the tyrosinase substrate L-DOPA, and spectrophotometry was used to quantify pigment production. Compared to both the wildtype and wildtype/heterozygote mix, *prdm1a -/-* mutants had significantly decreased pigment production (p< 0.01) and this loss of tyrosinase activity aligns with the levels of *prdm1a*, as lysates from a combination of wildtype and *prdm1a+ /-* embryos exhibited an intermediate loss of tyrosine activity that was significantly different from that of both the wildtype and *prdm1a-/-* lysates (**Figure 3G**). Together with our RT-qPCR results demonstrating that *prdm1a* mutants are deficient in the expression of late melanocyte markers, these data suggest that melanocyte progenitors require Prdm1a to terminally differentiate and produce pigment. Moreover, this suggests that even a partial loss of *prdm1a* affects melanogenesis, which is important to note given that our adult melanoma models are *prdm1a+ /-* heterozygotes. Together, the data indicate that Prdm1a is important for melanocyte development from neural crest cells. Given that there are key similarities between melanoma and neural crest cells and that PRDM1 is already a known tumor suppressor, we determined whether it played a role in melanoma as well.

### Low *PRDM1* expression is correlated with worse patient outcomes and metastatic melanoma in publicly available datasets

To determine whether *PRDM1* expression was associated with human melanoma patient prognosis, we analyzed The Cancer Genome Atlas (TCGA) cutaneous melanoma data set and plotted the overall survival of patients with metastatic melanoma based on their expression of *PRDM1*. Lower *PRDM1* expression is correlated with a worse overall survival rate (p< 0.001) (**Figure 4A**). We also mined the Riker melanoma data set (GSE7553 Moffitt data) and observed a significant decrease in *PRDM1* expression in metastatic melanoma tumors compared to normal skin (p< 0.001) and primary melanoma tumors (p< 0.05) (**Figure 4B**).

**Figure 4:**
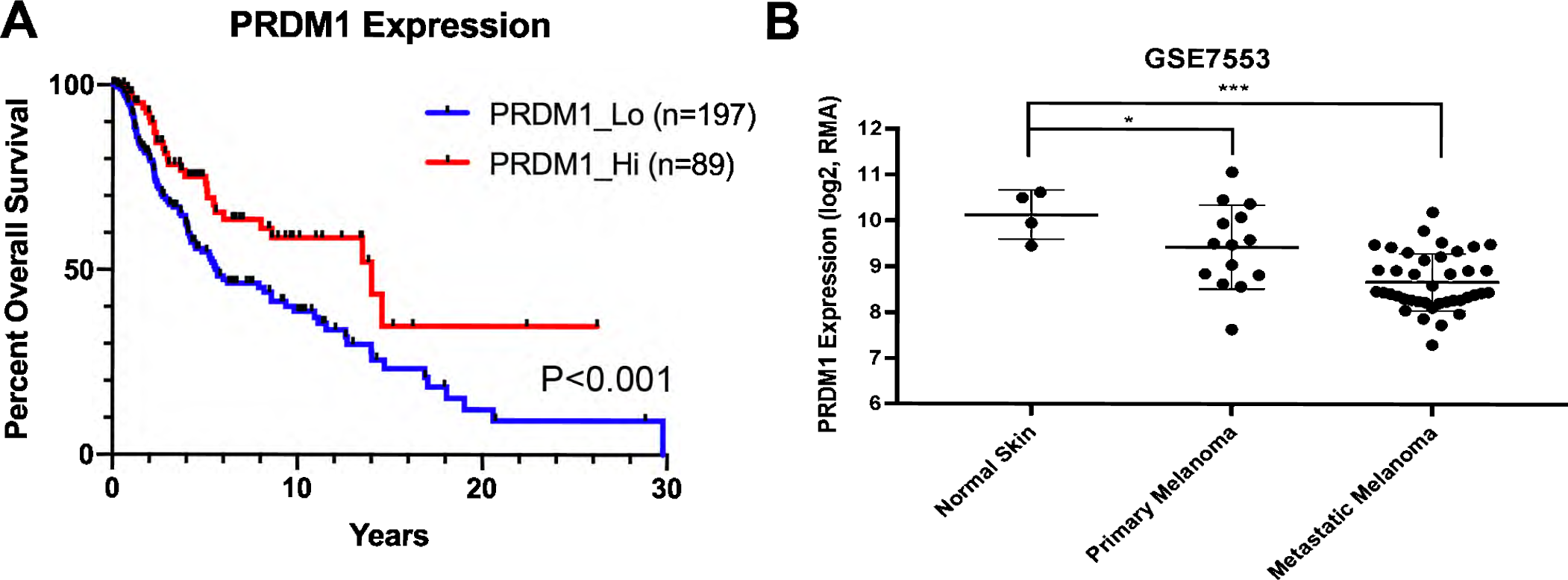
Low *PRDM1* expression in humans is correlated with worse patient outcomes and metastatic melanoma. (A) Overall survival of patients with metastatic melanoma compared to *PRDM1* expression obtained from The Cancer Genome Atlas (TCGA). Statistical analysis was performed with a log-rank test. (B) *PRDM1* expression in normal skin, primary melanoma tumors and metastatic melanoma tumors obtained from the Riker melanoma data set. Statistical analysis was performed using a one-way ANOVA. Error bars represent mean ± SD. *p< 0.05, **p< 0.01, ***p< 0.001

### Heterozygous loss of *prdm1a* in zebrafish increases melanoma onset and progression

To determine whether loss of PRDM1 promotes melanoma *in vivo*, we used two zebrafish models, an injected and transgenic *mitfa:*BRAF^V600E^ line. In both models, the BRAF mutation is driven by a *mitfa* promoter to drive melanoma in melanocytes specifically, with an additional *p53-/-* mutant background that further accelerates melanoma onset^17,40^. For the injected model, the oncogenic *mitfa*:BRAF^V600E^ construct was introduced into *p53-/-* and *p53-/-; prdm1a*+ /- zebrafish embryos at the single-cell stage^17,40^. Again, homozygous *prdm1a-/-* mutants die at 6-7 dpf. The construct is integrated randomly into the genome and is expressed transiently. We monitored the zebrafish and noted when hyperpigmentation occurred in concentrated areas of the body. This hyperpigmentation was indicative of the initiation of melanoma tumor formation. Due to transient expression of the *mitfa*:BRAF^V600E^ mutation in the injected model, tumor formation was slower than previously reported^17,20,40^. The *p53-/-* zebrafish formed tumors 12 months post-injection (n=12), while the *p53-/-;prdm1a+ /-* zebrafish developed tumors significantly more quickly at 10 months post-injection (n=8) (p< 0.05) (**Figure 4A**). Unfortunately, due to the *p53-/-* mutant background, the zebrafish often developed malignant peripheral nerve sheath tumors (MPNST)^59^ at about 10 months post-injection before melanoma formation. Therefore, these fish were excluded from our melanoma study.

To circumvent this issue, we established two stable transgenic lines, Tg(*mitfa*:BRAF^V600E^)*p53-/-* and Tg(*mitfa*:BRAF^V600E^)*p53-/-;prdm1a*+ /-, and monitored tumor formation. As expected, these tumors formed more quickly than in the injected model. By 17 weeks, about 50%of the zebrafish for both genotypes had formed tumors (**Figure 5B**). With one copy of *prdm1a* mutated, the Tg(*mitfa*:BRAF^V600E^);*p53-/-;prdm1a*+ /- zebrafish (n=33) quickly surpassed their wildtype Tg(*mitfa*:BRAF^V600E^);*p53-/-* siblings (n=18) and developed melanoma tumors (p< 0.05) (**Figure 5B**). By week 19, more than 60%of the Tg(*mitfa*:BRAF^V600E^);*p53-/-; prdm1a*+ /- zebrafish had visible tumors. In contrast, approximately 50%of the wildtype siblings were still tumor free, suggesting that loss of *prdm1a* accelerates tumor onset. Representative images of melanoma tumors are shown at 7 and 11 months for both genotypes (**Figure 5C**), and tumor area was measured and normalized to the body length (**Figure 5D**). The tumor area of Tg(*mitfa*:BRAF^V600E^);*p53-/-;prdm1a*+ /- (n=7) was significantly larger than wildtype Tg(*mitfa*:BRAF^V600E^);*p53-/-* tumors (n=5) (p< 0.05) (**Figure 5C-D**). However, this may be because tumor onset occurred earlier in Tg(*mitfa*:BRAF^V600E^);*p53-/-;prdm1a*+ /- zebrafish (**Figure 5B**). Similar to the injected model, several transgenic fish developed MPNST tumors due to the *p53-/-* mutant background^59^. Therefore, the melanoma tumors on these fish were not large enough to measure and were not included in the quantification.

**Figure 5:**
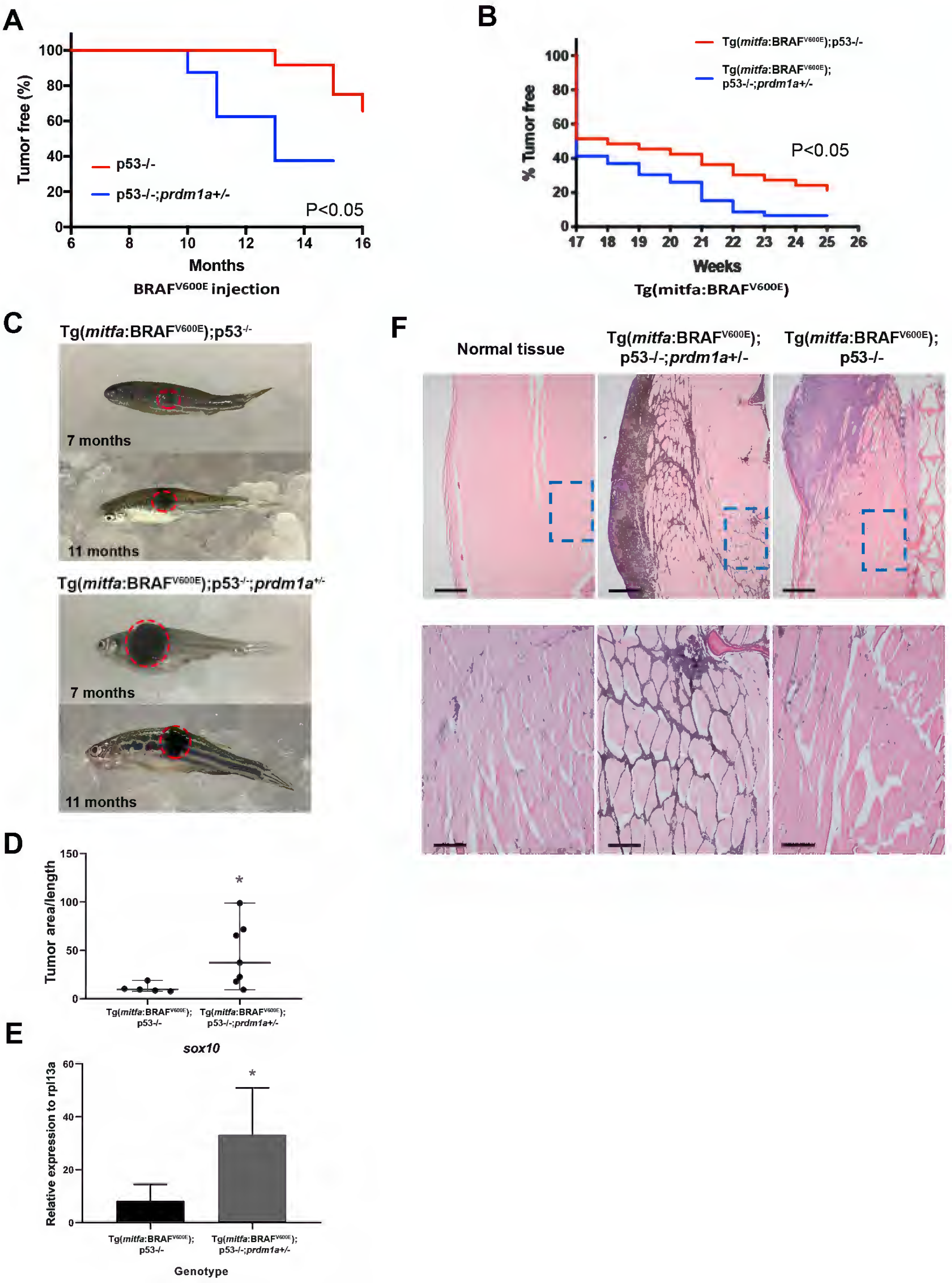
Loss of *prdm1a* in zebrafish accelerates tumor onset. (A) Tumor free curve from p53-/- (n=12) and p53-/-;*prdm1a+ /-* (n=8) zebrafish injected with BRAFV600E construct to induce melanoma. (B) Tumor free curve from Tg(*mitfa*:BRAF^V600E^);p53-/- (n=18) and Tg(*mitfa*:BRAF^V600E^);p53-/-;*prdm1a+ /-* (n=33) zebrafish. Statistical analysis for tumor free curves was performed with a log-rank test. (C) Representative images of melanoma tumors from Tg(*mitfa*:BRAF^V600E^);p53-/- and Tg(*mitfa*:BRAF^V600E^);p53-/-;*prdm1a+ /-* zebrafish at 7 and 11 months. (D) Tumor area was measured in five Tg(*mitfa*:BRAF^V600E^);p53-/- tumors and seven Tg(*mitfa*:BRAF^V600E^);p53-/-;*prdm1a+ /-* zebrafish tumors. The area was then divided by the body length to normalize. Relative tumor area was compared using an unpaired, Mann-Whitney U test. Error bars represent median and range. (E) RNA was isolated from dissected tumors and RT- qPCR was performed for *sox10.* Expression was compared using an unpaired, Mann-Whitney U test. Error bars represent mean ± SD. *p< 0.05. (F) H&E staining of size-matched tumors for Tg(*mitfa*:BRAF^V600E^);p53-/-;*prdm1a+ /-* (7 months) and Tg(*mitfa*:BRAF^V600E^);p53-/- (9 months) zebrafish compared to normal skin. Upper panels show 40X magnification. The lower panels are zoomed in to 200X magnification.

As previously described, Sox10 is a conserved marker of neural crest cells and potentially co-opted in melanoma progenitor cells in many vertebrate models^20,60^, thus we compared its expression between dissected Tg(*mitfa*:BRAF^V600E^);*p53-/-;prdm1a*+ /- and Tg(*mitfa*:BRAF^V600E^);*p53-/-* tumors. For each tumor, *sox10* expression was compared to normal, adjacent muscle tissue of the same individual. *sox10* expression varied but was elevated for both genotypes. On average, tumor samples from Tg(*mitfa*:BRAF^V600E^);p53-/-;*prdm1a*+ /- zebrafish had a threefold increase in *sox10* compared to their Tg(*mitfa*:BRAF^V600E^);*p53-/-* siblings (p< 0.01) (**Figure 5E**). This suggests that reduction of *prdm1a* expression causes an upregulation of *sox10* in both neural crest development and melanoma cells^21,61^. In melanoma, we predict that the gain in *sox10* expression supports the ability of melanoma to grow and spread.

To examine the zebrafish tumors in more detail, we paired zebrafish with similarly sized tumors and then sectioned the samples for H&E staining. In both tumors, we observed invasion into the neighboring muscle tissue with variable levels of melanin across the tumors (n=3 for each genotype; **Figure 5F**). The melanoma in Tg(*mitfa*:BRAF^V600E^);*p53-/-;prdm1a*+ /- zebrafish have an overall larger tumor area (**Figure 5D**), with melanin observed in the neighboring tissue (**Figure 5F**) whereas the Tg(*mitfa*:BRAF^V600E^);*p53-/-* tumor remains somewhat more localized at the skin surface with less invasion into the surrounding skeletal muscle.

## DISCUSSION

Melanoma cells often acquire features of neural crest cells during tumor formation by reactivating genes and pathways utilized by neural crest cells^12,13^. The work presented in this study suggest that Prdm1a, a known neural crest regulator, is required for melanocyte differentiation from neural crest cells and that loss of Prdm1a accelerates melanoma onset and progression. During development, we have shown that *prdm1a-/-* mutants have a reduction in neural crest derivatives, including mature pigment cells^25,26^. Here, we demonstrate that loss of *prdm1a* leads to an increase in expression of *sox10*, an early melanocyte stem cell marker, and an overall decrease in later melanocyte markers. *prdm1a-/-* mutants also have significantly reduced pigment production by tyrosinase. Together, these data suggest that Prdm1a is required for the differentiation of neural crest cells to mature melanocytes. It is likely that in the absence of Prdm1a, the cells are maintained as early melanocyte progenitors and thus function in supporting melanoma formation and progression. The case for *PRDM1* functioning as a tumor suppressor in melanoma is further supported by the fact that *PRDM1/prdm1a-c* copy number is significantly reduced in both human and zebrafish melanomas, and its expression is reduced in melanoma cells compared to normal melanocytes. Moreover, in human patients, low PRDM1 expression is correlated with worse human patient survival. Finally, we demonstrate that heterozygous loss of *prdm1a* promotes melanoma tumorigenesis in a zebrafish model and results in an increase in *sox10* expression in the tumors. Thus, our data demonstrates a novel role for the neural crest regulator PRDM1 as a melanoma tumor suppressor.

Melanoma is known to have increased somatic mutation rates compared to other cancers due to mutagenesis by UV exposure^49,50^, with *BRAF*^*V600E*^ being the most common mutation^3^. Other mutations, such as *NRAS*, are common in large congenital nevi, which are often melanogenic and can be precursors to melanoma^62^. Interestingly when *NRAS*^*Q61K*^ is driven by a tyrosinase promoter, transgenic mice develop congenital naevi and melanoma and express the neural crest progenitor marker Sox10^18^, again iterating that Sox10 expression is required for malignant melanoma formation. Reducing one copy of Sox10 in this background reduces the number of congenital naevi and melanoma tumors by regulating cell proliferation^18^. This suggests that Sox10’s function in development, to maintain neural crest stem cells, may be dysregulated during disease progression.

In our previous work and this current body of work, we have demonstrated that the developmental regulator, Prdm1a, is required for neural crest differentiation to mature melanocytes, which is consistent with its function in other cell types during development^29-31^. We show that *prdm1a* may promote differentiation through regulation of *sox10*^61^, as suggested by the increase in its expression in *prdm1a-/-* mutants. However, because the *prdm1a -/-* mutants have visibly less pigmentation compared to wildtype embryos, this may have affected our interpretation of the melanocyte marker gene expression data. Moreover, just from their reduced pigmentation and reduced expression of late melanocyte markers, we cannot conclusively claim that *prdm1a-/-* larvae in fact have less melanoblasts or melanocytes, as *prdm1a* loss may disrupt neural crest cells’ ability to differentiate into these cell types or impair the proliferation of these neural crest derivatives. In future studies, we can address these issues by performing live-imaging experiments and/or lineage tracing experiments to parse out the exact effects of *prdm1a* loss on neural crest derivatives. However, it is interesting that reducing *prdm1a* in heterozygous embryos also reduces tyrosinase activity, even when there is not a discernable difference in pigmentation. Therefore, our results suggest that Prdm1a is an important regulator of neural crest cell differentiation into melanocytes. Our data contribute to the growing literature regarding the parallels between neural crest cell development and melanoma tumor formation^13^.

Interestingly, PRDM1 is known to act as a tumor suppressor in other cancers, particularly B, T, and natural killer cell lymphomas and lung cancer^34,36,37,54^. We have now shown in a zebrafish model that heterozygous loss of *prdm1a* can accelerate melanoma onset, and it is likely acting as a tumor suppressor. We hypothesize that loss of *PRDM1* coordinates with mutations in *BRAF* and/or *TP53* to promote tumorigenesis. *PRDM1* deletions have been reported to be associated with loss of *TP53* in anaplastic large cell lymphoma patients^37^, and this association likely contributed to the pathogenicity of the cancer. Our zebrafish models carried mutations in both *BRAF* and *TP53*. Additional mutations in *PRDM1* led to accelerated melanoma tumor onset and suggested a trend towards metastasis (data not shown), similar to what has been shown with PRDM1 and lung cancer^54^. Whether *PRDM1* coordinates with these or additional tumor suppressor/oncogenic genes during tumorigenesis must be explored further.

In conclusion, this study demonstrates a novel role for PRDM1 as a tumor suppressor in *p53* mutated melanoma. We showed that even partial loss of *prdm1a* in zebrafish accelerates melanoma onset, and this can potentially be sourced to the impairment of melanocyte differentiation from neural crest cells during embryonic development observed upon *prdm1a* loss. Broadly, these results compound on accumulating evidence that neural crest cells and melanomas operate on overlapping gene regulatory networks and suggest that factors that regulate neural crest development are a source of therapeutic value for melanomas.

## Author Contributions

R.I. and B.T.T. contributed equally to this work, performed, analyzed, and interpreted data, and performed the statistical analysis on the zebrafish; A-C.T. performed bioinformatic analysis; D.O. performed histological analysis and imaging. K.B.A. conceptualized the project and obtained funding, collaborating with Y.S.; B.T.T. and K.B.A. wrote the manuscript with the help of Y.S.; and R.I., J.H., C.C. and A-C.T. edited the manuscript.

## Acknowledgments

We thank the Artinger and Shellman labs for experimental feedback; Morgan Singleton and UCD-AMC CLAC for zebrafish care; Virginia Ware and Aaron Clark for technical help with experiments; and Richard White for sharing microarray data, fish lines and advice.

The State of Colorado, Tobacco Settlement Funds pilot grant

SDRC Gates Center Pilot Grant, University of Colorado P30AR057212-10

The National Institutes of Health P30CA046934, University of Colorado Cancer Center Support Grant

## URLs

http://www.cbioportal.org/

http://cancergenome.nih.gov/

http://www.ncbi.nlm.nih.gov/gds\

